# Susceptibility to auditory closed-loop stimulation of sleep slow oscillations changes with age

**DOI:** 10.1101/2019.12.15.876847

**Authors:** Jules Schneider, Penelope A. Lewis, Dominik Koester, Jan Born, Hong-Viet V. Ngo

**Author notes:** **Corresponding authors:** Hong-Viet V. Ngo, Donders Institute for Brain, Cognition and Behaviour, Radboud University Medical Centre, Kapittelweg 29, 6525 EN Nijmegen, NL, Penelope A. Lewis.

## Abstract

**Background:** Cortical slow oscillations (SOs) and thalamo-cortical sleep spindles hallmark slow wave sleep and facilitate sleep-dependent memory consolidation. Experiments utilising auditory closed-loop stimulation to enhance these oscillations have shown great potential in young and older subjects. However, the magnitude of responses has yet to be compared between these age groups.

**Objective:** We examined the possibility of enhancing SOs and performance on different memory tasks in a healthy older population using auditory closed-loop stimulation and contrast effects to a young adult cohort.

**Methods:** In a within-subject design, subjects (n = 17, 55.7 ± 1.0 years, 9 female) received auditory click stimulation in synchrony with SO up-states, which was compared to a no-stimulation sham condition. Overnight memory consolidation was assessed for declarative word-pairs and procedural finger-tapping skill. Post-sleep encoding capabilities were tested with a picture recognition task. Electrophysiological effects of stimulation were compared to those reported previously in a younger cohort (n = 11, 24.2 ± 0.9 years, 8 female).

**Results:** Overnight retention and post-sleep encoding performance of the older cohort revealed no beneficial effect of stimulation, which contrasts with the enhancing effect the same stimulation protocol had in our younger cohort. Auditory stimulation prolonged endogenous SO trains and induced sleep spindles phase-locked to SO up-states in the older population. However, responses were markedly reduced compared to younger subjects. Additionally, the temporal dynamics of stimulation effects on SOs and spindles differed between age groups.

**Conclusions:** Our findings suggest that the susceptibility to auditory stimulation during sleep drastically changes with age and reveal the difficulties of translating a functional protocol from younger to older populations.

**Highlights:**

- Auditory closed-loop stimulation induced SOs and sleep spindles in older subjects
- Stimulation effects were reduced and overall susceptibility diminished with age
- Slow oscillation and sleep spindle dynamics deviated from those in younger subjects
- Stimulation shows no evidence for memory effect in older subjects

## 1. Introduction

Sleep is an integral part of our lives, and research has accumulated strong evidence regarding its importance for maintaining health [1]. In particular slow wave sleep (SWS) supports crucial immunological, endocrine, metabolic, and cognitive functions [2–6]. For instance, the intricate interplay of its hallmark <1 Hz slow oscillations (SOs) and 12-15 Hz fast spindles facilitates sleep-dependent memory consolidation by orchestrating the transfer of newly acquired information into long-term storage and the strengthening of memory traces [7]. Likewise, the ability to acquire novel information is contingent on priorly obtained SWS [8–11].

Sleep quality naturally declines during both healthy and pathological ageing [12]. Linked to age-related neural atrophy, the number and amplitude of SOs decreases, and overall power in the 0.5-4 Hz slow wave band is strongly reduced [13–17]. Moreover, decline in SWS during later-life brain ageing predicts deterioration of memory abilities [18]. In older adults, the benefit of sleep for consolidation is either reduced or non-existent [19–22]. Furthermore, shallower sleep is associated with less successful encoding of novel information post-sleep in healthy older individuals [10,23].

Experiments to counter such impairments by enhancing sleep are highly topical and have been successfully trialled with several stimulation modalities [24–28]. Auditory closed-loop stimulation has proven to be a particularly promising technique [29], which consists of detecting endogenous SOs and applying brief auditory stimuli during their positive peaks. This method has been shown to induce both SO and fast spindle activity, thereby boosting performance on a declarative memory task in young adults overnight [29–33]. However, mixed effects were found in healthy older adults [34,35], and the extent to which stimulation effects depend on age and beneficially impact on other forms of memory in older subjects are presently unknown. In the current study, we investigated whether an auditory closed-loop stimulation targeting SOs in a group of older adults would likewise enhance sleep oscillations, declarative and procedural memory consolidation, as well as post-sleep encoding abilities. Furthermore, by drawing on a previously reported dataset of healthy young adults who showed substantial memory benefits from the same stimulation protocol [29], we directly compared physiological stimulation effects between these two age groups.

## 2. Materials & methods

### 2.1 Subjects

Seventeen healthy volunteers (9 female) aged 49 to 63 years (mean ± SEM = 55.7 ± 1.0) with no history of psychological, neurological, or sleep disorders were recruited. All were non-smoking, native German speakers and had followed a regular sleep/wake schedule for four weeks prior to participation. Subjects were screened for good hearing (3-digit hearing test) and no signs of mild cognitive impairment (score ≥ 24/30 in the Montreal Cognitive Assessment), nor excessive daytime sleepiness (Epworth Sleepiness Scale, mean ± SEM score = 10.83 ± 0.91). The experiment received ethical approval from the Universities of Tübingen and Manchester. All subjects gave informed written consent before participation.

In order to compare electrophysiological effects of auditory closed-loop stimulation in our older cohort to a younger population, we took advantage of a previously published dataset [29]. This consists of 11 healthy young adults (8 female, mean ± SEM age = 24.2 ± 0.9 years), who fulfilled matching participation requirements and underwent identical stimulation procedures.

### 2.2 Experimental design and procedure

An initial adaptation night accustomed subjects to sleeping in the sleep laboratory, and was followed by at least one recovery night at home. In a within-subject design, subjects then spent two counterbalanced experimental nights in the laboratory (Fig. 1A), undergoing one experimental stimulation (Stim) and one control condition without stimulation (Sham) each and at least 7 nights apart. On experimental days, subjects were instructed to wake at 7 am, not consume any alcohol within the prior 24 hours, and not ingest caffeine after 2 pm. Upon arrival at the sleep laboratory at 8 pm, subjects were prepared for polysomnography. They then performed a psychomotor vigilance task (PVT), followed by a declarative paired associate learning (PAL) and procedural finger-tapping (FT) task. Prior to bedtime at ~11 pm, they completed the Stanford Sleepiness Scale (SSS). Subjects were awakened the following morning while in stage 1 or 2 NREM sleep after ~8 hours of sleep opportunity. At least half an hour after waking, subjects completed a questionnaire to assess subjective sleep quality [36] and the SSS, followed by PAL recall, a post-sleep-only picture encoding (PE) recognition task interleaved with a distractor Digit Span Task, and lastly an FT retest. See supplementary materials for details on memory and control tasks.

**Figure 1:**
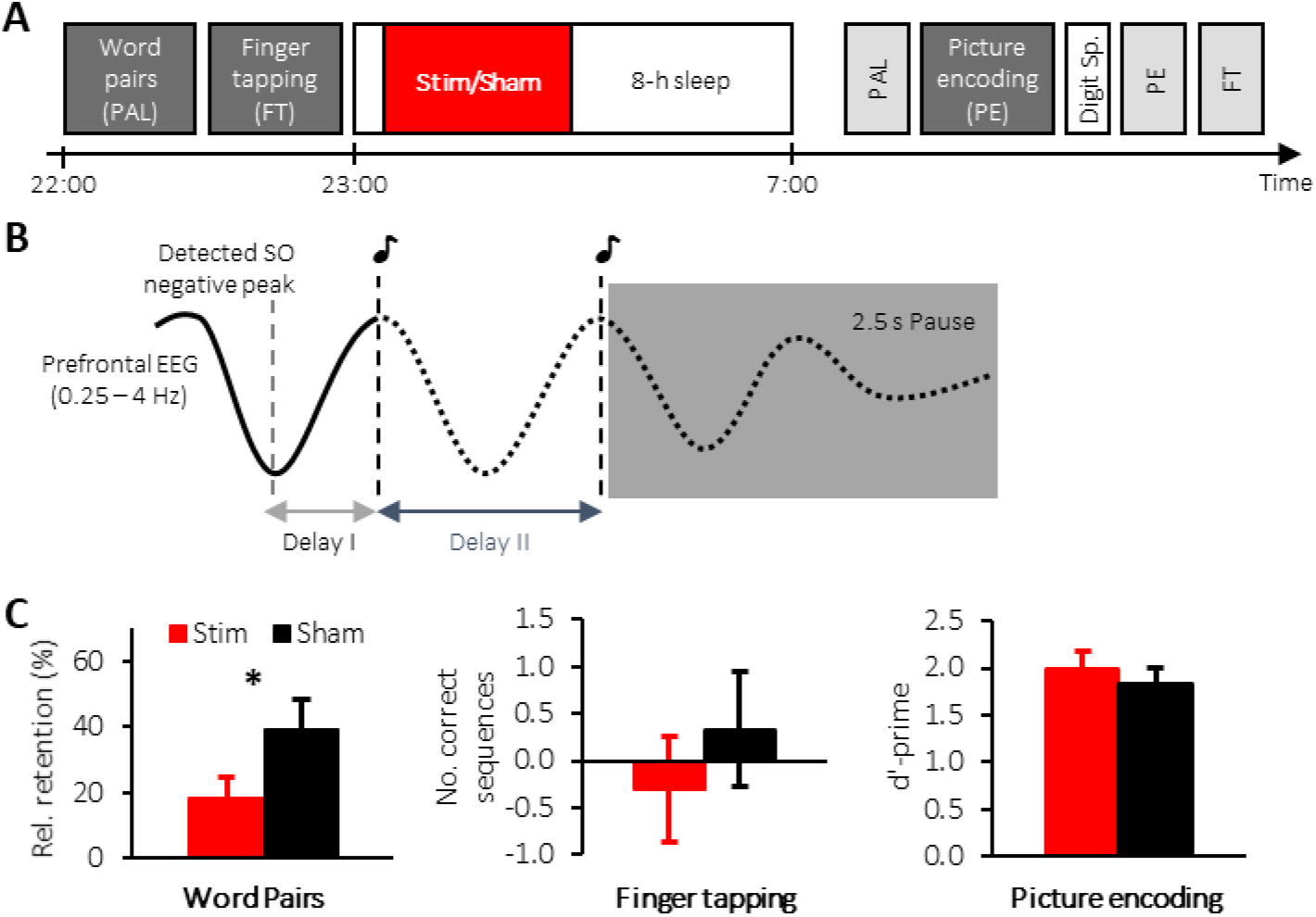
Study design and behavioural results for older population. **(A)** Subjects learned word-pairs (PAL) including an immediate cued recall and were then trained on a finger tapping (FT) task. Afterwards subjects were allowed to sleep for 8-hours during which (in the first 180 min) auditory closed-loop stimulation (Stim) or no stimulation (Sham) was applied. The next morning, word-pair memory (PAL) was tested, followed by a picture encoding (PE) task, for which the encoding and recognition phases were interleaved by a Digit Span Task. Finally, finger tapping (FT) performance was tested. **(B)** Schematic illustrating the stimulation protocol. Upon detection of an SO negative peak, the first and second click were delivered after two individually adapted delays (Delays I and II) followed by a stimulation pause of 2.5 s. **(C)** Mean ± SEM of memory performance on the PAL (left), FT (middle) and PE (right) tasks for the Stimulation (red) and Sham condition.

### 2.4 Polysomnography recordings

Polysomnography was continuously recorded with a BrainAmp DC amplifier (Brain Products, Gilching, Germany) with scalp electroencephalogram (EEG) electrodes positioned according to the international 10-20 system at F3, Fz, F4, C3, Cz, C4, P3, Pz, P4, all referenced to linked mastoids. One ground electrode was placed on the forehead. Impedances were kept below 5 kΩ. EEG data were sampled at 500 Hz and saved on a computer for later analyses. In addition to the above setup, an additional six electrodes were attached to record horizontal and vertical electrooculography, and electromyography for standard polysomnography.

### 2.5 Auditory closed-loop stimulation

Real-time identification of spontaneous SOs was based on a previously described algorithm [29] (see supplementary materials for details). Upon detection of a SO down-state, a first auditory click was presented after an individually determined delay (delay I: mean ± SEM = 583.24 ± 26.50 ms) to coincide with the subsequent SO up-state (Fig. 1B). A second click was delivered after a second individual delay (delay II: mean ± SEM = 1091.47 ± 21.06 ms) concurring with the upcoming induced SO up-state, followed by a detection pause of 2.5 s. Delay II was introduced due to the variant nature of SO parameters in this age group and adjusted in eleven of 17 older subjects, unlike in the original study where this was kept constant at 1075 ms [29]. Stimulation commenced once subjects had spent ~5 min in stable NREM sleep and was continued for 3.5 hrs, but stopped manually for arousals or changes in sleep stage. In Sham nights, an identical protocol was followed, i.e. SO up-states were identified but no click sounds delivered. Stimuli consisted of 50 ms pink 1/f noise and were delivered binaurally via earbuds (Sony MDR-EX35) at individually adjusted volume levels, which had been previously determined during SWS in the adaptation night (mean ± SEM volume = 54.53 ± 1.17 dB). Upon questioning in the mornings following experimental nights, only three individuals reported hearing the sounds.

### 2.6 EEG analysis

All data analyses were carried out in Matlab (Version R2016b, MathWorks, USA) and the Fieldtrip toolbox [37] using custom-made scripts. Following an initial filtering of EEG and EOG between 0.3-30 Hz, and of EMG above 5 Hz, two trained experimenters determined sleep stages across experimental nights according to standard scoring criteria [38] while blinded to experimental condition. Sleep stages S1, S2, SWS (= S3 + S4), REM sleep, wake, and epochs containing movement and other arousals were identified from lights off until waking time. Percentage of time spent in each stage was calculated as time in the respective sleep stage over total sleep time (TST).

To calculate event-related auditory responses, the EEG signal during SWS-epochs was averaged in windows of 5 s, with a 2 s pre-stimulus offset with regard to the first stimulus. Analysis of evoked fast spindle activity followed the same procedure with an additional bandpass filtering between 12-15 Hz, calculation of the root mean squared signal (RMS, based on a window of 200 ms), and a baseline correction between −2 to −1.5 s.

To assess spatiotemporal patterns of evoked responses (measured by the large negative component at ~500 ms post-stimulus) and the fast spindle response, we examined the relative change of the induced responses with respect to the endogenous (initially detected) SO for the stimulation condition. To this end, we divided the largest negative amplitude value found 0 to 1 s post-stimulus for the first and second click respectively by the mean baseline value of the endogenous SO between −1 to 0 s preceding the first click (Fig. 2C). Contrarily, for the fast spindle response, we determined the ratio of the largest peak in the fast spindle RMS signal between 0.5 and 1.5 s after each stimulus to a baseline value obtained between −0.5 and 0.5 s centred around the first click (Fig. 3C). Furthermore, to evaluate the impact on spindle refractoriness, we calculated the difference in the mean fast spindle RMS activity derived from a pre-stimulus interval between −2 and −1.5 s and late post-stimulus interval between 2.5 and 3 s with respect to the 1st click (Fig. 3D).

**Figure 2:**
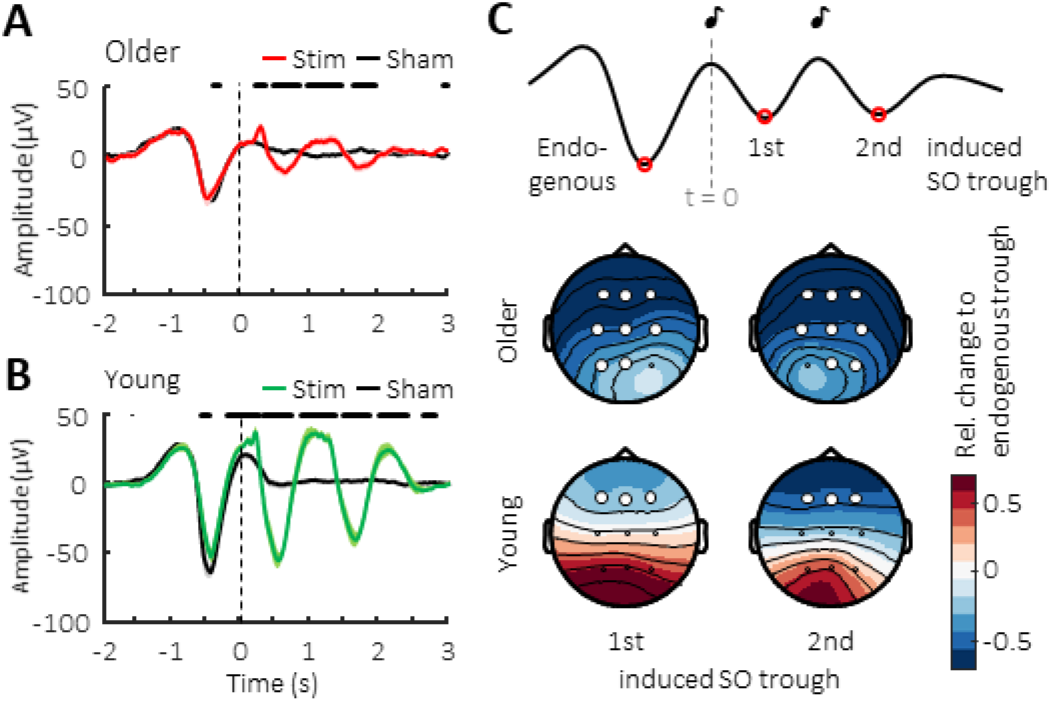
Event-related potentials upon auditory click stimulation. **(A)** Mean ± SEM EEG-signal from Cz averaged time-locked to the first click for the Stimulation (red) and Sham conditions (black) in older population. Vertical line indicates timing of the first clicks, whereas thick horizontal black bars mark time points of significant difference between conditions. **(B)** Mean ± SEM EEG-signal from Cz averaged time-locked to the first click for the Stimulation (green) and Sham conditions (black) in young cohort. Vertical line indicates timing of the first clicks, whereas thick horizontal black lines at the top mark time points of significant difference between conditions. **(C)** Top schematic illustrates the time points during which trough amplitudes were obtained to determine the relative change shown colour-coded as topographical maps of the evoked response with respect to the endogenous SO. Vertical grey line marks time point of the first click (t = 0). White circles indicate channel location with a significant change from baseline after FDR correction.

**Figure 3:**
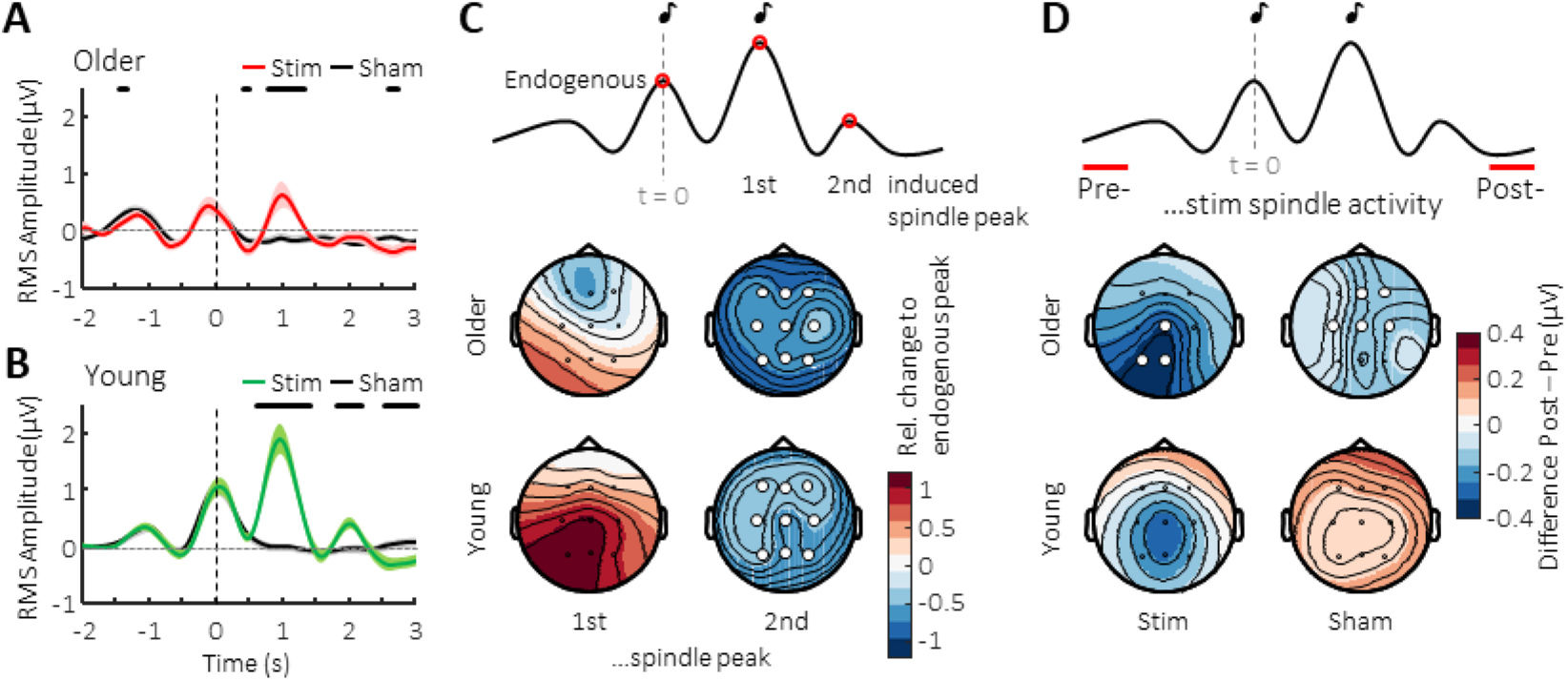
Immediate effects on fast spindle activity. Mean ± SEM RMS-signal in the 12-15 Hz spindle-band from Cz averaged time-locked to the first click for Sham (black) and Stimulation conditions in **(A)** the older population (red) and **(B)** the young adult group (green). Vertical line indicates timing of the first click, whereas thick horizontal black bars mark time points of significant difference between conditions. **(C)** Top schematic illustrates the time points during which fast spindle peak activity was obtained to determine the relative change of the evoked response with respect to the endogenous SO shown below colour-coded in topographical maps. **(D)** Topographic distribution of the colour-coded difference in RMS-spindle activity between two 500-ms intervals preceding (Pre) and following (Post) acute two-click stimulation or Sham-trials for the older and younger population, as illustrated in the schematic above. Vertical grey line marks time point of the first click (t = 0). White circles indicate channel location with a significant relative change **(C)** or difference **(D)** from baseline after FDR correction.

Sustained stimulation effects were examined by spectral analysis during SWS using Fast Fourier Transformation (based on 8.2 s segments with 50% overlap and a Hanning window) across the 3.5-h stimulation period. SO peak frequency between 0.5-1.25 Hz as well as the mean power in the 12-15 Hz fast spindle range were then determined. SO peak frequency was chosen as a measure to fully capture the displayed SO frequency variance in the older cohort. Moreover, we detected discrete SOs during SWS epochs across the entire night based on previously described algorithms [39,40] (see supplementary materials for details). The number of offline detected SOs was then determined for the ~3.5 h stimulation period. Furthermore, for the same period, we calculated the mean SO peak-to-peak amplitude, and phase-locked fast spindle activity was examined by averaging the fast spindle RMS-signal time-locked to the SO trough of detected events in a window from −1.25 to 1.25 s with a baseline correction from −1.25 to −1.15 s. Finally, to assess the temporal interrelationship among SOs during the ~3.5 h stimulation period, we examined for each offline-detected SO event at electrode Cz the occurrence of pre- and succeeding SOs based on event histograms within 100 ms bins and in a ± 3 s time interval around the trough (at t = 0). Resulting histograms were normalised by the total number of detected SO events (multiplied by 100) and then the difference between Stim – Sham conditions was calculated.

### 2.7 Statistical analysis

All data are shown as mean ± SEM. Statistical analyses were generally based on paired-samples student’s t-tests, or repeated-measures analyses of variance (RM ANOVA) with possible within-subject factors ‘condition’ (Stimulation vs. Sham), ‘SO trough or spindle peak’ (1st and 2nd induced response), ‘topography’ (9 EEG channels), and between-subject factor ‘age’ (young vs. older adults). We focus on reporting significant interactions and age group main effects. Separate post-hoc ANOVAs were subsequently conducted for each age group. If necessary, a Greenhouse-Geisser correction for degrees of freedom was applied. Topographical plots were prepared based on channel-wise paired-tests with adjustment for multiple comparison using false discovery rate (FDR) corrections [41]. The threshold of significance was set to a *p value* < 0.05. In the case of missing individual data values due to technical error (n = 1 in PVT) or subjects omitting questionnaire items (n = 2 in sleep quality), the respective individuals were excluded from corresponding analyses.

## 3. Results

### 3.1 Memory performance in older adults is unaffected by stimulation

We first examined whether stimulation affected overnight retention across different memory tasks in the older cohort. Contrary to expectations, baseline normalised declarative word pair memory was worse following a night of stimulation (18.68±6.22% vs. 39.48±8.76% for the Stimulation and Sham condition, respectively, with t(16) = −2.13, P = 0.049, Fig. 1C, left). Moreover, stimulation had no impact on overnight change in procedural finger-tapping skill (change in number of correctly tapped sequences, Stimulation: −0.30±0.56, Sham: 0.33±0.62; t(16) = −1.01, P = 0.328, Fig. 1C, middle), and also did not affect post-sleep encoding of pictures (d’ = 2.04±0.19 and d’ = 1.85±0.17 for Stimulation and Sham condition, t(16) = 1.62, P = 0.125, Fig. 1C, right).

### 3.2. Evoked responses in the older cohort are weaker than in younger subjects

We next set out to examine the electrophysiological responses to the stimulation and made use of an existing EEG dataset from a young adult cohort (see section 2.1 for details), allowing for direct contrasting between the two age groups. Averaging the EEG signal time-locked to the first stimulus revealed that the SO rhythm was prolonged by two additional cycles in older subjects (Fig. 2A). However, a comparison to the younger cohort (Fig. 2B) illustrates that these immediate responses are immensely diminished with age. A direct comparison of the stimulation-induced effects relative to the pattern observed in Sham demonstrated markedly greater enhancement in amplitudes in the young adults compared to the older cohort (Supplementary Fig. 1A). To quantify this observation, we determined within each cohort the difference in amplitude between the detected endogenous SO trough and the evoked troughs following both the first and second click in the stimulation condition only (Fig. 2C). This revealed a clear difference in trough amplitudes between age groups (F(1,26) = 5.91, P = 0.022 for the age-group × SO trough interaction). A follow-up examination for each age group showed no difference between the amplitudes of endogenous and elicited SO troughs in young adults (F(1,10) < 0.747, P > 0.408 for main effects of SO trough). By contrast, the older group exhibited significantly smaller elicited amplitudes compared to the endogenous trough, with a relative decrease down to approximately 50% of the endogenous SO amplitude (main effects of SO trough: F(1,16) < 62.01, P < 0.001). This pattern of diminished response in the older but not young adults indicates a difference in susceptibility to auditory stimulation during sleep in the older population.

### 3.3 Older subjects exhibit stronger fast spindle refractoriness

Following the notion that fast sleep spindles co-occur with SO up-states, we examined responses in the 12-15 Hz fast spindle frequency range. We extracted spindle RMS activity and averaged the signal time-locked to the first auditory stimulation to confirm an increase in spindle power phase-locked to the first induced up-state in older adults (Fig. 3A). This was absent from the second induced up-state (at ~2 s post-stimulus), mirroring the pattern previously observed in young adults (Fig. 3B). However, similar to the overall evoked response shown earlier, the initial increase in fast spindle power was diminished in the older population when directly compared to the younger cohort (see Supplementary Fig. 1B). To determine the difference between endogenous and elicited fast spindle responses in both age groups, we next calculated the difference between the endogenous fast spindle peak, i.e. at the time of the first click presentation, and the induced fast spindle peaks (~1 and 2 s post-stimulus) within each cohort in the stimulation condition. This analysis first indicated an overall difference between age groups (main effect ‘age group’ with F(1,26) = 4.46, P = 0.044), and, secondly, a difference in change in topography across induced spindle peaks (F(1.99, 51.75) = 3.81, P = 0.029 for the spindle peak x topography interaction). A decomposition into the two age groups to explore the topographic pattern of the first induced response (compared to the baseline peak) revealed a similar response strength during the endogenous and first induced spindle peak in the older cohort (F(1,16) = 0.017, P = 0.899, Fig. 3C). In the younger group by comparison, this induced spindle response was almost twice the size of the preceding endogenous baseline peak (F(1,10) = 6.87, P = 0.026).

With regard to the absence of a spindle response following the 2nd click, the older subjects exhibit a pattern similar to the young group in terms of a strong subsequent suppression of fast spindle power. However, a closer visual inspection of the older cohort (Fig. 3A) suggests that such suppression is also present in the Sham condition and may continue for a longer duration after the first SO than in younger adults. To examine these potential differences in spindle refractoriness induced by the stimulation between age groups in more detail, we contrasted spindle RMS activity obtained before (t = −2 to −1.5 s, “pre”) and after (t = 2.5 to 3 s, “post”) acute double-click stimulation (Fig. 3D), and found a significant difference in its spatiotemporal pattern (F(2.26,38.47) = 5.10, P = 0.008 for the interaction of spindle window (pre vs. post x topography x age)). A consecutive decomposition into separate age groups showed that while our young subjects did not exhibit spindle suppression in the post time window in either condition (Stimulation: F(1,10) = 2.64, P = 0.135, Sham: F(1,10) = 1.25, P = 0.291), the older population showed a decrease in fast spindle power during the late time window in both Stimulation and Sham conditions (F(1,16) = 7.81, P = 0.013 and F(1,16) = 13.96, P = 0.002 for Stimulation and Sham, respectively). This pattern suggests that besides an overall change in susceptibility to auditory stimulation during sleep, the dynamics of spindle-expressing thalamo-cortical networks are altered in the ageing brain, with stronger refractory periods observed in older compared to young adults.

### 3.4 Opposite overall effects of stimulation on SOs and fast spindles between age groups

In order to assess the overall influence of the stimulation irrespective of click presentations, we next turned our attention to spectral power and also identified discrete SO events post-hoc within the entire stimulation period.

For spectral SO peak power, we found a striking difference between the cohorts (F(1,26) = 17.67, P < 0.001 for the age group x condition interaction). This effect was in particular mediated by an overall stimulation-related increase in SO peak power in the younger subjects (main effect for condition with F(1,10) = 22.73, P < 0.001, Fig. 4A), but not the older subjects (F(1,16) = 0.99 for condition main-effect, P = 0.334). Meanwhile, an identical spectral analysis tailored to the 12-15 Hz fast spindle band returned no significant effects of stimulation or age differences (P ≥ 0.262 for all main and interaction effects).

**Figure 4:**
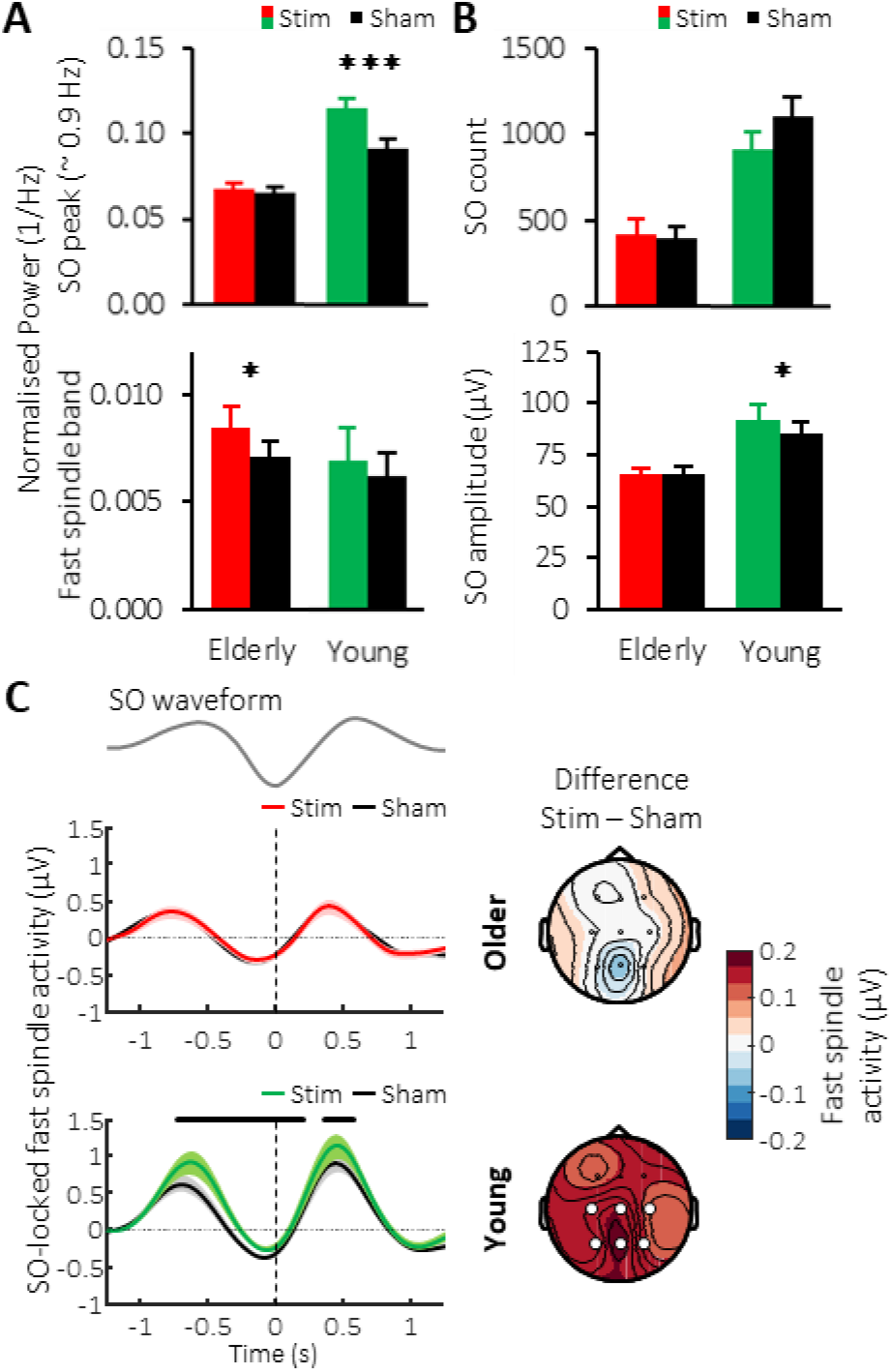
Sustained modulation of SOs and fast spindles. **(A)** Global mean ± SEM of the normalised spectral power for the SO peak obtained across the 180-min stimulation period for the Stimulation condition in the older (red) and young population (green) and their corresponding Sham conditions. **(B)** Mean ± SEM of SO amplitude of offline-detected SO events across the stimulation period for the stimulation condition in the older (red) and young (green) cohort and their Sham conditions (black). **(C)** Fast spindle RMS-activity average time-locked to the negative peak (vertical lines) of offline detected SO events for the older (top) and young population (below) with Stimulation conditions shown in red or green, and Sham conditions in black. Thick horizontal black bars mark time points of significant difference between conditions. The corresponding topographical distribution of the difference between conditions over the time intervals −1.25 to 1.25 s (where t = 0 time-locked to the negative SO peak) is shown on the right. White circles indicating channel locations with significant difference in overall phase-locked fast spindle activity between conditions after FDR correction.

Initial examination of the number of discrete SO events occurring during the stimulation period confirmed an overall lower number of SO events in the older population (F(1,26) = 27.95, P < 0.001). However, neither group showed any difference in SO numbers between experimental conditions (Young: F(1,10) = 3.17, P = 0.105, Older: F(1,16) = 0.46, P = 0.508). Instead, we found a modulation of the amplitude of discrete SO events (F(1,26) = 8.68, P = 0.007 for the age x condition interaction), especially driven by an increased amplitude difference between the Stimulation and Sham conditions in the younger participants (Fig. 4B). No differences were observed between groups or conditions for fast spindle power or SO count across the stimulation period (Supplementary Fig. 1C top and bottom). Interestingly, examining the occurrence of SO events with event-histograms revealed no sustained prolonging of SO trains by the stimulation in older compared to the young adults (Supplementary Fig. 1D). Together, these results confirm a non-resonant SO response in the ageing brain.

Finally, we investigated the co-occurrence of SO and sleep spindles during the stimulation period, given their hypothesized joint contribution to memory consolidation. Averaging the fast spindle RMS signal time-locked to the SO down-state confirmed increases in fast spindle activity during SO up-states in both cohorts and conditions. However, relative to the corresponding Sham condition, stimulation caused a significant difference in overall grouped fast spindle activity between groups (F(1,26) = 7.05 and P = 0.013 for the Group x Condition interaction), which was driven by a widespread elevation of fast spindle activity during the SO up-to-down transition in young adults (F(1,10) = 8.10, P = 0.017), but not in the older cohort (F(1,16) = 0.06, P = 0.808, Fig. 4C).

### 3.5 Closed-loop stimulation in older age does not alter sleep architecture and other control measures

Table 1 contains the general sleep parameters for the older group (please see Supplementary Table 1 for young subjects). As previously reported for the young subjects [29], analyses demonstrated that auditory stimulation was not associated with any change in sleep onset (t(16) = −0.17, P = 0.867), total sleep time (t(16) = 0.55, P = 0.588), or overall sleep architecture for the stimulation period (all P ≥ 0.193) or the entire night (all P ≥ 0.266) in older adults. Auditory stimulation did not result in disrupted sleep through increased arousals (t(16) = −1.27, P = 0.216). Moreover, sleep questionnaires and control tasks revealed no influence of auditory stimulation on subjective sleep measures or control tasks (all P ≥ 0.103, see Table 2 for an overview).

**Table 1:** Sleep architecture during the 3-hour stimulation period and entire night in the older adults. Stimulation did not alter time spent in any of the sleep stages, total sleeping time, or number of arousals. TST = total sleep time, S1-S2: sleep stages 1 and 2, SWS = Slow wave sleep (i.e. S3 + S4), REM = rapid eye movement.

|  | Stim |  |  | Sham |  |  | P-value |
| --- | --- | --- | --- | --- | --- | --- | --- |
|  | mean | ± | SEM | mean | ± | SEM |  |
| **TST (min)** | 449.32 | ± | 10.05 | 443.85 | ± | 13.56 | 0.588 |
| **Sleep onset (min)** | 10.91 | ± | 2.31 | 11.26 | ± | 2.13 | 0.867 |
| **Stimulation period** |  |  |  |  |  |  |  |
| **Wake (%)** | 9.75 | ± | 3.15 | 11.07 | ± | 3.15 | 0.552 |
| **S1 (%)** | 3.55 | ± | 0.59 | 3.84 | ± | 0.60 | 0.623 |
| **S2 (%)** | 56.95 | ± | 3.64 | 60.57 | ± | 3.27 | 0.242 |
| **SWS (%)** | 13.31 | ± | 2.91 | 11.85 | ± | 2.46 | 0.462 |
| **REM (%)** | 16.25 | ± | 1.86 | 12.48 | ± | 2.13 | 0.193 |
| **Arousal index (%)** | 7.01 | ± | 0.64 | 8.17 | ± | 1.18 | 0.388 |
| **Entire Night** |  |  |  |  |  |  |  |
| **Wake (%)** | 12.07 | ± | 2.97 | 9.83 | ± | 1.70 | 0.423 |
| **S1 (%)** | 5.17 | ± | 0.54 | 5.67 | ± | 0.54 | 0.375 |
| **S2 (%)** | 56.08 | ± | 2.89 | 59.13 | ± | 2.10 | 0.266 |
| **SWS (%)** | 8.17 | ± | 1.64 | 7.38 | ± | 1.69 | 0.508 |
| **REM (%)** | 17.89 | ± | 1.52 | 17.68 | ± | 1.94 | 0.912 |
| **Arousal index (%)** | 6.64 | ± | 0.77 | 8.22 | ± | 0.91 | 0.216 |

**Table 2:**
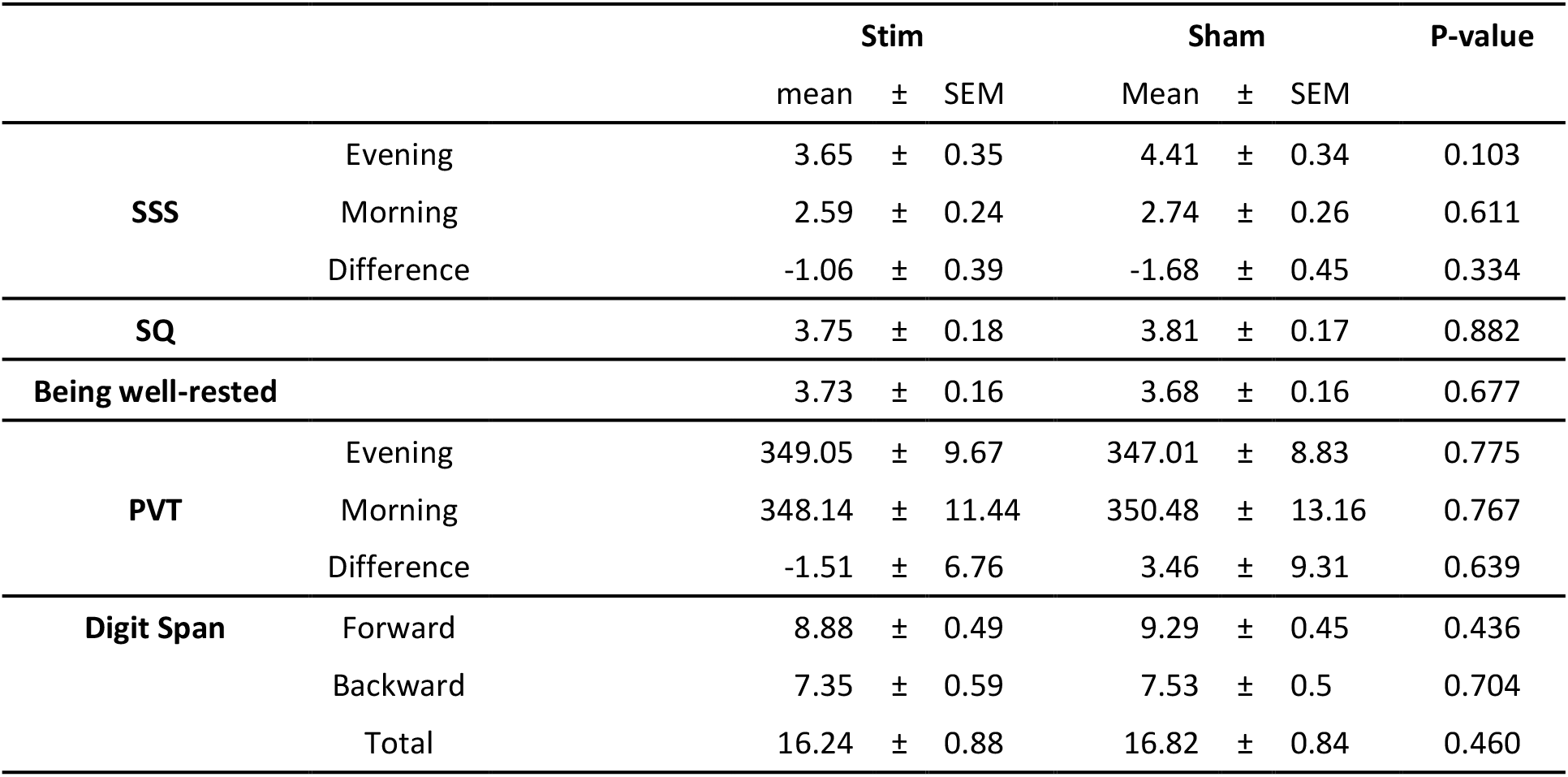
Subjective measures and control tasks. Stimulation did not impact on Stanford Sleepiness Scale (SSS) ratings, subjectively reported sleep quality (SQ) and feelings of being well-rested, or on performance on the psychomotor vigilance (PVT) or Digit Span tasks.

## 4. Discussion

Our data confirm the feasibility of selectively inducing SOs in older subjects using auditory closed-loop stimulation. However, stimulation did not promote performance on any of the assessed memory domains in the older cohort, but rather impaired the retention of declarative memories. Brain responses of older adults were quantitatively diminished and revealed different patterns for SOs and fast spindles in comparison to a younger population, indicating a change in susceptibility to stimulation with age.

The ageing brain shows a distinct physiological response to clicks administered in a closed-loop manner and an absence of apparent sleep detriments, i.e. increased arousals elicited by the sound clicks, as might be expected in shallower and more fragmented sleep. However, when contrasting physiological enhancement between conditions in young and older adults, the extent to which SO trough amplitude could be enhanced by stimulation in relation to a preceding, endogenous SO trough was approximately halved in older adults. The older cohort also did not demonstrate sustained increases in SO measures, e.g. in amplitude as observed in young adults, suggesting stimulation effects to be more short-lived by comparison. Potential reasons for this decreased susceptibility of the older brain to stimulation could range from smaller engaged cortical populations and un-timely cortical reactivity of neurons, to decreased connectivity preventing cardinal sleep rhythms from longer-term resonance, possibly due to prolonged cell refractoriness. A combination of these factors could prevent the ageing brain from responding to incoming stimuli as strongly as a younger brain [42,43]. Whether altered SO characteristics in older age hence require different stimulation settings, i.e. adapted timings for stimuli to occur at a particular phase ‘sweet spot’ where greater enhancement could still be elicited, remains a subject for future investigation.

In keeping with original findings in the young cohort, stimulation compared to baseline in older subjects led to an increase in fast spindle activity on the first, but not second induced SO peak [32]. Similarly to the reduced strength observed in overall evoked responses, the power of both endogenous and induced fast spindles in our older group was roughly half of the value measured in young adults. Moreover, whereas fast spindle refractoriness was evident during stimulation in both age groups, signs of stronger spindle refractoriness across subsequent SO cycles were also observed in the Sham condition in older adults. This implies altered thalamo-cortical network dynamics in the ageing brain, which, once a spindle has been expressed, require more time to recover and re-establish baseline levels in cellular reactivity. Thus, age-dependent changes in spindle-expressing networks, i.e. increased spindle refractoriness even during unstimulated conditions, appear to present a critical physiological limitation to stimulation enhancement. Our analysis on the interplay of SOs and fast spindles indicated topographically widespread stimulation-induced increases in fast spindle activity nesting within SO up-states in the young adult group only. By contrast, this spindle increase was entirely missing in the older cohort. Interestingly, the older group did not show a decoupling between SO and spindles, i.e. no systematic shift of the preferential phase of sleep spindles occurring within SOs as has recently been reported in other studies as a factor critically contributing to the sleep induced enhancement in memory [44,45].

The short-lived increases in SO power and unaltered SO-locked fast spindle activity in the older cohort provide a conceivable explanation for the stimulation’s unfavourable impact on memory performance in this group. Similar to other studies [35], stimulation did not improve performance on the declarative task in the older adults. On the contrary, stimulation even impaired memory retention. Recent studies revealed opposing effects on memory for SOs and delta waves, strengthening and weakening memories, respectively [46,47]. Against this backdrop, stimulation in older adults in the present study may have preferentially enhanced delta waves characterized mainly by distinctly smaller (down state) trough amplitudes in comparison with SOs. However, post-sleep encoding was likewise unaffected by stimulation, presumably due to a lack of increase in SO power during the stimulation period to effectively re-establish hippocampal retention capacity overnight, similar to [33]. Since a predefined amplitude threshold must be crossed to trigger stimulation, a decline in endogenous SO number, density, and amplitude in older age [15–17] naturally results in fewer stimulation opportunities [48]. Adjusting the detection threshold may prove useful in such cases. Papalambros and colleagues [34] found a positive impact of auditory closed-loop stimulation on declarative memory performance when boosting SWA in older individuals. However, their phase-locked loop algorithm worked with a threshold which, at −40 μV, was set on average only half as high as ours and may have targeted oscillations which strictly speaking no longer qualify as SOs based on the amplitude criterion [49]. Additionally, their experiment administered stimulation throughout the entire night. These factors likely resulted in a proportionally larger fraction of stimulated endogenous SOs.

To conclude, the present study demonstrated that auditory closed-loop stimulation can be applied to the ageing brain without being detrimental to sleep architecture. However, stimulation outcome diverged considerably between older and younger subjects. Our results suggest the magnitude and nature of inducible enhancement is reduced in the ageing brain, a pattern reported in previous studies using different stimulation techniques [48,50,51]. The inability to influence sleep-dependent memory consolidation is most likely due to a combination of age-related changed characteristics of cardinal sleep rhythms, and physiological and cellular constraints, as observed in SO generation and fast spindle-expressing network dynamics. Despite this decreased susceptibility to auditory closed-loop stimulation in older age, the fact that similar stimulation efforts [34] have yielded positive behavioural results emphasises the challenges of translating a functional protocol to different age groups. It is all the more evident that future research is necessary to elucidate the nature of these cardinal sleep rhythms and their functions in order to aid the development of real-world clinical applications, e.g. to counter decline in healthy and pathological ageing.

## Acknowledgements

The authors would like to thank Wael El-Deredy, Jason Taylor, and Gareth Gaskell for valuable discussions, Neil Pendleton for sharing his gerontological expertise, and all subjects for their participation. JS, PL, JB & HVVN designed the experiment; JS, DK & HVVN performed the experiment; JS & HVVN analysed the data, and JS, PL, JB & HVVN interpreted results and wrote the paper.

