## Supplementary Materials and Methods for "Susceptibility to auditory closed-loop stimulation of sleep slow oscillations changes with age"

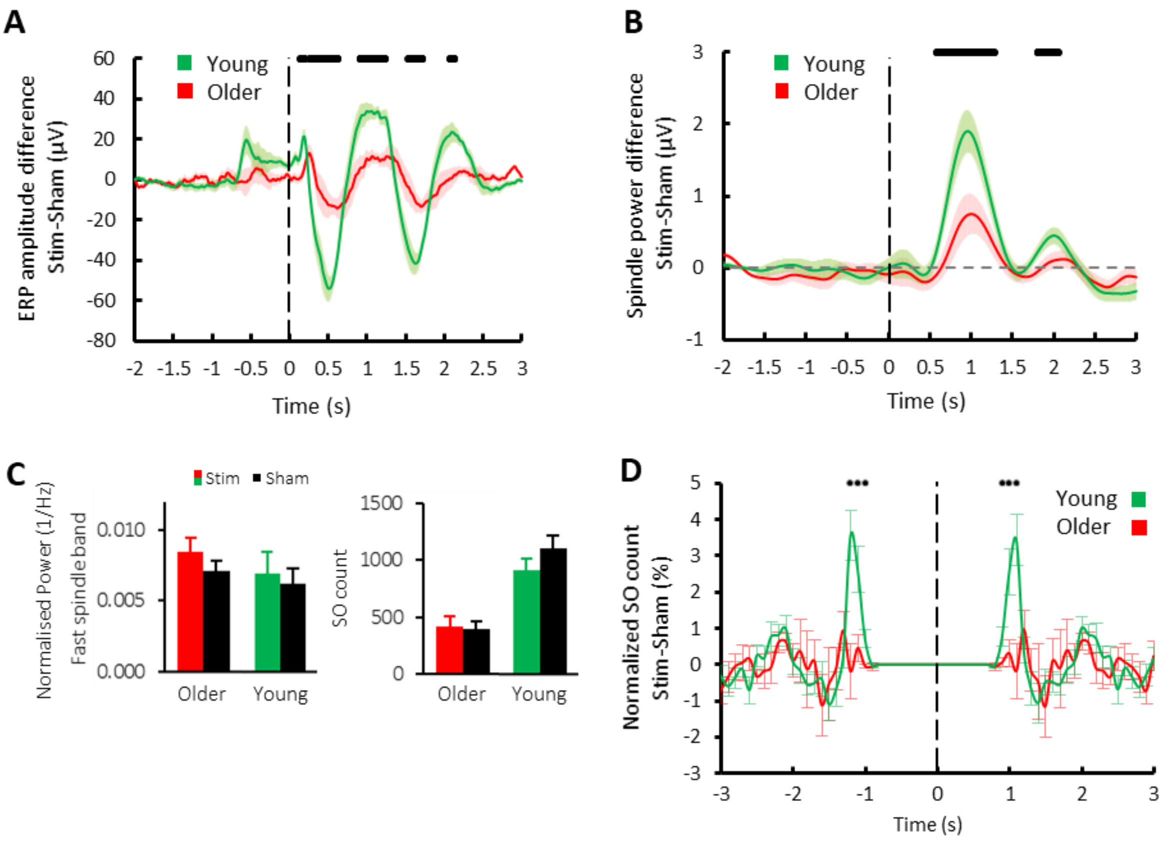


**Supplementary Figure 1: (A)** Comparison of event-related potentials upon auditory stimulation between groups. Mean ± SEM EEG-signal from Cz averaged time-locked to the first click for Stimulation - Sham conditions in older (red) and young (green) populations. **(B)** Group differences in immediate stimulation-induced fast spindle RMS effects (Stim - Sham). Mean ± SEM RMS-signal in the 12-15 Hz spindle-band from Cz averaged time-locked to the first click in the older population (red) and the young adult group (green). Vertical line indicates timing of the first click, whereas thick horizontal black bars mark time points of significant difference between groups. **(C)** Global mean ± SEM of the normalised spectral power for the fast spindle frequency band obtained across the stimulation period for the stimulation condition in the older (red) and young population (green) and their corresponding Sham conditions (top) and SO count of offline-detected SO events across the stimulation period (bottom) for the stimulation condition in the older (red) and young (green) cohort and their Sham conditions (black). **(D)** Auto-event histogram of offline-detected SO events reveals no sustained prolonging of SO trains in older subjects. To assess the temporal interrelationship among SOs during the ~3 h stimulation period, we examined for each offline-detected SO event the occurrence of pre- and succeeding SOs based on event histograms within a ±3 s time interval and 100 ms bins. Resulting histograms were normalised by the total number of detected SO events (multiplied by 100) and then the difference between Stim – Sham conditions was calculated. Mean ±

SEM for young (green) and older adults (red) are pictured at representative electrode Cz. Time t = 0 (vertical black line) denotes the negative peak of detected SO events. Black dots denote statistically significant differences between the young and older cohorts (uncorrected).

| | Stim | | | Sham | | | P-value |
|---|---|---|---|---|---|---|---|
| | mean | ± | SEM | mean | ± | SEM | |
| **TST (min)** | 413.5 | ± | 6.6 | 417.6 | ± | 2.8 | 0.490 |
| **Sleep onset (min)** | 10.9 | ± | 5.4 | 12.4 | ± | 4.0 | 0.810 |
| **Stimulation period** | | | | | | | |
| **Wake (%)** | 6.3 | ± | 1.6 | 7.3 | ± | 1.1 | 0.545 |
| **S1 (%)** | 8.2 | ± | 1.0 | 7.5 | ± | 1.4 | 0.662 |
| **S2 (%)** | 48.6 | ± | 2.7 | 47.3 | ± | 2.2 | 0.676 |
| **SWS (%)** | 29.1 | ± | 4.1 | 28.7 | ± | 2.8 | 0.905 |
| **REM (%)** | 7.7 | ± | 1.6 | 9.2 | ± | 1.8 | 0.148 |
| **Arousal index (%)** | 6.7 | ± | 0.4 | 6.7 | ± | 1.0 | 0.883 |
| **Entire Night** | | | | | | | |
| **Wake (%)** | 5.8 | ± | 1.2 | 5.4 | ± | 0.9 | 0.639 |
| **S1 (%)** | 9.6 | ± | 1.0 | 7.0 | ± | 0.8 | 0.014 |
| **S2 (%)** | 49.0 | ± | 2.6 | 50.3 | ± | 1.7 | 0.587 |
| **SWS (%)** | 20.1 | ± | 2.4 | 19.0 | ± | 2.0 | 0.606 |
| **REM (%)** | 15.5 | ± | 1.2 | 18.3 | ± | 1.8 | 0.061 |
| **Arousal index (%)** | 7.9 | ± | 0.6 | 6.7 | ± | 0.5 | 0.212 |

**Supplementary Table 1: Sleep architecture during the 3-hour stimulation period and entire night in the young adult cohort.** Stimulation did not alter time spent in any of the sleep stages (except for total night S1), total sleeping time, or number of arousals. TST = total sleep time, S1-S2: sleep stages 1 and 2, SWS = Slow wave sleep (i.e. S3 + S4), REM = rapid eye movement.

**Memory and control tasks**

*Paired associate learning task.* To assess declarative memory, subjects were instructed to learn 80 word pairs of semantically moderately related German nouns [29]. Different sets of pairs were used in counterbalanced lists between subjects and nights, with different word pair order in all learning and recall sessions. In evening sessions, subjects were asked to carefully study the word pairs when each was presented for 4 s on screen with an interstimulus interval of 1 s. A subsequent immediate recall test established baseline retention by presenting the first noun of the word pair and requiring recall of the second noun. Subjects were given unlimited time and received feedback of the correct response. The morning recall session followed the same procedure but included no feedback. Overnight retention was calculated as the difference between the number of correct pairs obtained in the morning and evening divided by the evening performance (relative change).

*Finger tapping task.* Procedural memory was assessed on a finger-tapping sequence task [52]. Following a short practice round, subjects used the four fingers of their non-dominant hand to tap on a computer keyboard a fixed five-digit sequence presented on screen as often and accurately as possible within 30 s intervals. In the evening session, they completed 12 blocks of 30 s, interspaced with 30 s breaks. Feedback in the form of number of correct and overall attempted sequences was shown on screen following each block. The morning retest session followed an identical procedure only consisting of three blocks. Evening and morning performance scores were calculated by averaging the numbers of correctly tapped sequences in the last and first three blocks in the evening and morning sessions, respectively, with their difference representing overnight change.

*Picture-encoding task.* In the morning sessions only, a picture-encoding task presented subjects with 50 photographs of neutral landscapes and houses for 2.5 s each and then prompted them to indicate via keyboard presses whether the photo depicted a residential house or tropical landscape to aid encoding [8,10]. Picture presentation order was randomised, with a varying interstimulus interval of 0.6-1.4 s. Following the encoding phase, a ~5 min distractor Digit Span Task was conducted to distract subjects from mentally rehearsing stimuli between encoding and recall. To assess encoding performance, subjects were presented with 100 photographs, 50 of which they had previously been exposed to, and asked to indicate via keyboard presses whether they remembered previously seeing the picture, with answer options of ‘yes’, ‘maybe yes’, ‘maybe no’ and ‘no’. Encoding performance was evaluated by combining the former and latter two options respectively, counting cases of correctly remembered items (hit), incorrectly remembered items (false alarm), correctly negated items (correct rejection) and falsely negated items (miss). Accounting for response bias, we then calculated a final score d-prime (d’) by subtracting the z-transformed false alarm rate from the z-transformed hit rate.

*Digit span task.* Subjects were tasked to memorise and immediately verbally relay a number series of increasing length as read out by the experimenter. In the first part of the task and starting at level 1 with 2 trials of 3 digits, each further sequence increased in length by one digit per level up to level 7, but was ended whenever a subject repeated both trials per level incorrectly. Part two then required the subjects to repeat sequences backwards, with each corresponding level consisting of one less digit than the forward repeat part. Different sequences were used for each experimental night. Scores were calculated per forward, backward, and total number of trials and level reached.

*Psychomotor Vigilance Task.* A ~5 min psychomotor vigilance task (PVT, based on [53]) programmed in a custom-made software was used to measure subjects’ alertness and vigilance at the beginning of evening and morning sessions. Subjects were instructed to focus on a millisecond counter, which repeatedly appeared at the centre of the screen after a random delay ranging between 2-10 s, and to stop it with a key response as quickly as possible. An average score was calculated from all their responses with delay times below 150 ms and above 800 ms excluded. PVT data from one individual were lost due to a technical error.

**Setup and algorithm for real-time detection of SOs**

The present study used the same technical setup and stimulation protocol from [29]. A ’Digitimer D360’ EEG Amplifier (Digitimer, Welwyn Garden City, UK) and ‘Power1401 mk 2’ data acquisition interface (Cambridge Electronic Design, UK) were linked to a computer, all separate to the polysomnography. This facilitated real-time filtering between 0.25-4 Hz of the incoming EEG signal streaming from one additional second forehead ground and one EEG electrode placed on Fpz to allow a custom-designed algorithm in Spike2 (Cambridge Electronic Design, Cambridge, UK) to detect whenever the signal value dropped below a previously defined threshold, indicating an SO down-state during SWS (SO trough threshold was set to -80 µV by default, but had to be adjusted to -50 µV and -60 µV for one and two subjects, respectively). Furthermore, every 0.5s, the detection threshold was updated to the minimal instantaneous EEG amplitude within the preceding 5s interval, however, only if this value was smaller than the default threshold. The number of stimulation trials showed a between-subjects age group effect ($F(1,26) = 6.86$, $P = 0.015$), main effect for condition ($F(1,26) = 62.03$, $P < 0.001$) but no age x condition interaction ($F(1,26) = 0.26$, $P = 0.616$; mean ± SEM for older adults: Stim = 183.29±26.75, Sham = 401.82±41.90, & young adults: Stim = 320.27±45.03, Sham = 568.91±65.35).

**Offline detection of discrete SO events**

To identify discrete SO events offline each EEG channel was first high-pass filtered at 0.3 Hz (Butterworth 3rd order, two-pass) followed by a low-pass filtered at 1.25 Hz (Butterworth, 6th order, two-pass). Then positive to negative zero crossings were identified and all intervals between consecutive zero crossings shorter than 0.8 or longer than 2 s (corresponding to frequencies of 0.5–1.25 Hz) were discarded. Across the remaining intervals the negative peaks and the amplitude from negative to the positive peak were averaged. The resulting mean values were multiplied by 1.25 and served as detection threshold, i.e. intervals were labelled as a SO whenever its negative peak was lower than 1.25 times the mean negative peak value and the amplitude exceeded the 1.25 times the mean amplitude threshold. In order to perform SO-locked EEG-analysis, the negative trough was used a temporal reference point for individual SO events.
